# Revisiting neoantigen depletion signal in the untreated cancer genome

**DOI:** 10.1101/2020.05.11.089540

**Authors:** Shixiang Wang, Xuan Wang, Tao Wu, Zaoke He, Huimin Li, Xiaoqin Sun, Xue-Song Liu

## Abstract

This study is arising from Van den Eynden J. et al. Nature Genetics. Lack of detectable neoantigen depletion signals in the untreated cancer genome. Van den Eynden J. *et al.* tried to address a very important scientific question: could the immune system eliminate cancer cells with immunogenic mutations in untreated situation? Van den Eynden J. *et al.* first annotated the human exome into “HLA-binding regions” and “non HLA-binding regions” based on the predicted binding affinity of nonapeptides translated from the un-mutated reference coding genome with type I HLA alleles. They hypothesized that if neoantigen depletion signal exist, the nonsynonymous mutations in “HLA-binding regions” will be negatively selected during cancer evolution, while nonsynonymous mutation in “non HLA-binding regions” will not be negatively selected. This will lead to decreased nonsynonymous vs synonymous mutation ratio (n/s) in “HLA-binding regions” compared with “non HLA-binding regions”. They defined HLA-binding mutation ratio (HBMR) as the ratio of n/s in “HLA-binding regions” to “non HLA-binding regions”, and reported that HBMRs are close to 1 in different types of cancer after background corrections, meaning neoantigen depletion signals are not detectable in different types of cancer. The fundamental problem of their hypothesis lies in that the actual neoantigens with immunogenicity do not overlap with their defined “HLA-binding regions”. Actually, most neoantigens with immunogenicity are not located in “HLA-binding regions”, when dissimilarity between mutant and wild type peptide are considered. It is the neoantigen with immunogenicity, but not nonsynonymous mutation in their defined “HLA-binding regions” undergo immunoediting based negative selection. Thus the results reported in that study are fundamentally flawed, and at this current stage we could not draw a solid conclusion as to whether the neoantigen depletion signal exists or not.

## Main text

Immunoediting, which includes three temporally distinct stages, termed elimination, equilibrium, and escape, has been proposed to explain the complex relationship that exists between cancer cells and the immune system during the development of an overt malignancy ^1^. The interaction between tumor cells and the immune system undoubtly exists. However, to what extend these interactions sculpt the tumor genomic DNA alterations is still an open question. Recently Van den Eynden J. *et al.* reported that the neoantigen depletion signal is undetectable in the untreated cancer genome; however their method for detection of this neoantigen depletion signal is questionable as described below.

Van den Eynden J. *et al.* annotated “HLA-binding regions” in the human genome based on the predicted binding affinity of nonapeptides translated from the un-mutated reference coding genome with type I HLA alleles ^2^. Ratio of nonsynonymous vs synonymous mutation number (n/s) has been used as an internal indicator for evolutional selection ^3^. For example, nonsynonymous mutations in *TP53* are undergoing positive selection in cancer, and consequently *TP53* gene has high n/s value compared with background, while nonsynonymous mutations in *TTN* are not undergoing positive selection, and n/s value of *TTN* is comparable to background level.

Van den Eynden J. *et al.* hypothesized that if neoantigen depletion signal exists, nonsynonymous mutations should happens at a lower rate due to neoantigen depletion in their defined “HLA-binding regions” compared with non HLA-binding regions. Genomic alterations that encode neoantigen with sufficient immunogenicity can be selectively depleted due to immunoediting.

Van den Eynden J. *et al.* take it for granted that nonsynonymous mutations in their defined “HLA-binding regions” will encode neoantigen with immunogenicity, while in “non HLA-binding regions” nonsynonymous mutations will not encode neoantigen with immunogenicity. The fundamental problem of their hypothesis lies in that the actual neoantigens with sufficient immunogenicity does not overlap with their defined “HLA-binding regions” (Fig. 1a). It is real neoantigen with immunogenicity, but not nonsynonymous mutation in their defined “HLA-binding regions” undergoing immunoediting based negative selection. Thus the results reported in that study are fundamentally flawed, and at this current stage we could not draw a solid conclusion as to whether the neoantigen depletion signal exists or not.

**Fig.1.**
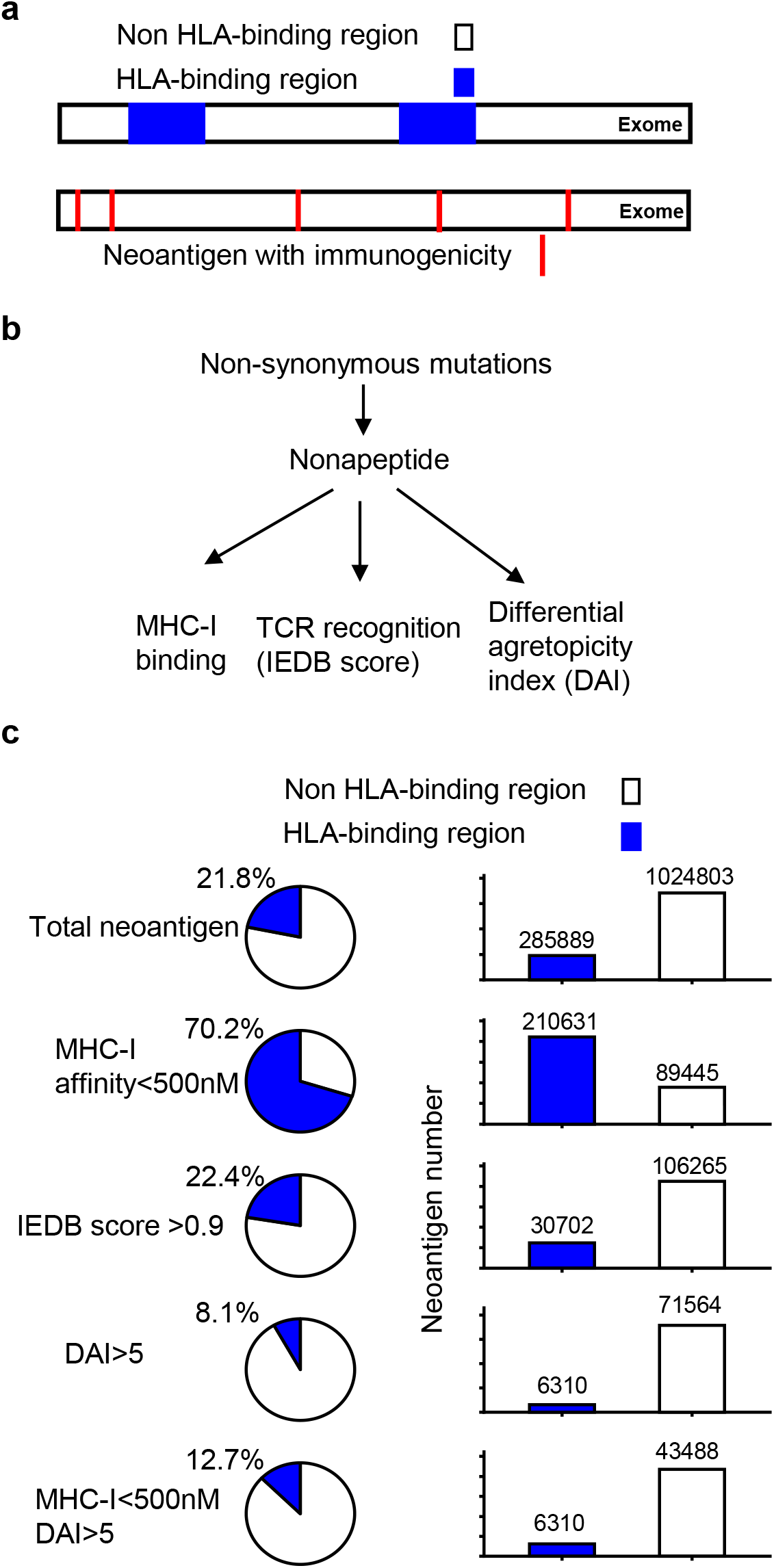
HLA-binding regions and neoantigens with immunogenicity. **a**, Schematic diagram about the “HLA-binding regions” and neoantigens with immunogenicity. “HLA-binding regions” were defined by Van den Eynden J. *et al.* as the un-mutated exome regions that code for nonapeptides with predicted binding affinity with type I HLA alleles less than 500nM. **b**, Workflow for identifying neoantigens with immunogenicity. **c**, Distributions of neoantigens with immunogenicity as defined by four different methods in “HLA-binding regions” and “non HLA-binding regions”.

It has become clear that tumor cells can evoke immune response, and have immunogenicity, and this is different from normal somatic cells, where tumor cells originally come from. The immunogenicity of tumor cells is determined by the genomic DNA alterations in tumor cells. How to define a neoantigen with sufficient immunogenicity is still not very clear. The golden standard to evaluate the immunogenicity of neoantigen relies on *in vivo* experiments. For example, test the immunogenicity of synthetic neopeptide with animal model, or detection of reactive immune cells against the neopeptide in clinical samples from cancer patients. Current methods to systematically evaluate the immunogenicity of neoantigen depend on *in silico* predictions with parameters including: binding with HLA I/II, TCR recognition, dissimilarity with un-mutated wild type counterpart ^4^.

We systematically evaluated the distribution of neoantigens with immunogenicity in human cancer genome. 1,836,369 mutations (525,677 synonymous and 1,310,692 nonsynonymous) from TCGA datasets located in the 17,992 genes with unambiguous protein information are available for analysis. 21.8% of the nonsynonymous mutations (285,889) are located in the “HLA-binding regions”, and this is consistent with their original report that “HLA-binding regions” occupy 22.1% of exome coding region. Neoantigens with immunogenicity were selected based on binding with MHC-I, TCR recognition, and dissimilarity with un-mutated wild type counterpart (Fig. 1b).

Initially, the selection of neoantigen with sufficient immunogenicity is based on the predicted MHC-I binding affinity < 500nM. In total, 300,076 neoantigens have a predicted MHC-I binding affinity < 500nM (Extended Data Fig. 1). Among these neoantigens, 210,631 (70.2%) are located in the “HLA-binding regions”, and 89,445 (29.8%) are not located in the “HLA-binding regions” (Fig. 1c). An enrichment of neoantigens with MHC-I binding affinity < 500nM in “HLA-binding regions” compared with non-HLA-binding region is observed (Odds ratio=29.27) (Extended Data Fig. 2), likely due to the contribution of un-mutated amino acid in MHC-I binding prediction.

TCR recognition probability was evaluated through the similarity analysis of tested neopeptide to known immunogenic epitopes in the immune epitope database (IEDB) ^5,6^. IEDB scores are in the range of 0-1, higher IEDB score means higher TCR recognition probability ^6^. In total 136,967 IEDB high neoantigens (IEDB score > 0.9) has been obtained (Extended Data Fig. 1). Among these IEDB high neoantigens, 30702 (22.4%) are located in the “HLA-binding regions”, and 106,265 (77.6%) are not located in the “HLA-binding regions” (Fig. 1c). No enrichment of IEDB high neoantigens in “HLA-binding regions” has been observed (Odds ratio=1.04) (Extended Data Fig. 2).

It is evident that the MHC binding score alone is not a robust predictor of immunogenicity or tumor rejection. Another important feature of neoantigen with sufficient immunogenicity is the distinction from wild type un-mutated peptide. In case when wild type counterpart peptide is immunogenic, T cells responsive to mutated neopeptide may have been centrally deleted or peripherally tolerated due to immunotolerance to wild type peptide ^7^. And several studies have already demonstrated that sequence dissimilarity is an important feature of immunogenicity ^6–10^. Differential agretopicity index (DAI), defined as the difference in binding strength between the mutated neopeptide and its un-mutated normal peptide counterpart has been widely used in prediction of neoantigen with immunogenicity ^7^. Higher DAI means more immunogenicity. Using DAI>5 as a criteria, in total 77874 neoantigens are obtained (Extended Data Fig. 1). Among them only 6310 (8.1%) are located in the “HLA-binding regions”, and 71,564 (91.9%) are not located in the “HLA-binding regions” (Fig. 1c). When both DAI>5 and MHC-I affinity <500nm are used, in total 49,798 neoantigens are obtained (Extended Data Fig. 1). Among them 6310 (12.7%) are located in the “HLA-binding regions”, and 43,488 (87.3%) are not located in the “HLA-binding regions” (Fig. 1c). We can clearly see from this analysis that most neoantigens with immunogenicity are not located in “HLA-binding regions”, when dissimilarity between mutant and wild type peptide are considered (Extended Data Fig. 2). This analysis questioned the rationale used by Van den Eynden J. *et al.* to detect the potential neoantigen depletion signal, and it is not unexpected that neoantigen depletion signal cannot be found in their defined “HLA-binding regions”.

This study does not question the model Van den Eynden J. *et al.* used for correcting background n/s rate, and in this respect their efforts are valuable for future studies. However their selection of “HLA-binding regions” is questionable, and their study cannot support their argument that neoantigen depletion signal is undetectable.

In summary, the key problem of Van den Eynden J. *et al.* study is that the actual genomic regions undergoing neoantigen depletion selection are not the “HLA-binding regions” defined in their study. When dissimilarity between mutant and wildtype peptide is considered, most neoantigens with immunogenicity are not located in their defined “HLA-binding regions”. In addition, some synonymous mutation could contribute to neopeptide generation through alternative splicing ^11^, and this situation can decrease the detection power of n/s ratio based methods. All together, based on their study we could not conclude if the neoantigen depletion signal exists or not. It is not unexpected that neoantigen depletion signal cannot be found in their defined “HLA-binding regions”.

## Methods

### Data download and statistical analysis

TCGA mutation data with aggregated HLA binding affinity score were downloaded from Van den Eynden J. *et al.* (zenodo https://doi.org/10.5281/zenodo.2621365). Differential agretopic index (DAI), defined as the ratio of binding strength between the mutated neopeptide and its unmutated normal peptide counterpart. Neoantigens were identified based on three approaches: 1) MHC affinity < 500nM; 2) differential agretopicity index (DAI) > 5; 3) IEDB score > 0.9. Overlaps between neoantigen locations and HLA binding/non-binding region from Van den Eynden J. et al. were calculated by BEDtools ^12^, then the results were analyzed and visualized by R 3.6. Fisher extract test was used to implement enrichment analysis.

## Data availability

Raw data has been provided in the supplementary files, other raw neoantigen list data are available upon request.

## Code availability

Custom R scripts used in this study are available at https://github.com/XSLiuLab/neoantigen_depletion.

## Acknowledgements

We thank the TCGA project for making cancer genomics data available for analysis. We thank Raymond Shuter for editing the text. Thank ShanghaiTech University High Performance Computing Public Service Platform for computing services. Thanks also to other members of Liu lab for helpful discussion.

## Contributions

S.W., X.W., T.W. performed research and analyzed data. Z.H., H.L., X.S. participated in critical discussion. X.-S.L. conceptualized this study and drafted the manuscript. S.W., X.W., T.W., X.-S.L. reviewed and edited the manuscript.

---

**Extended Data Fig.1.**
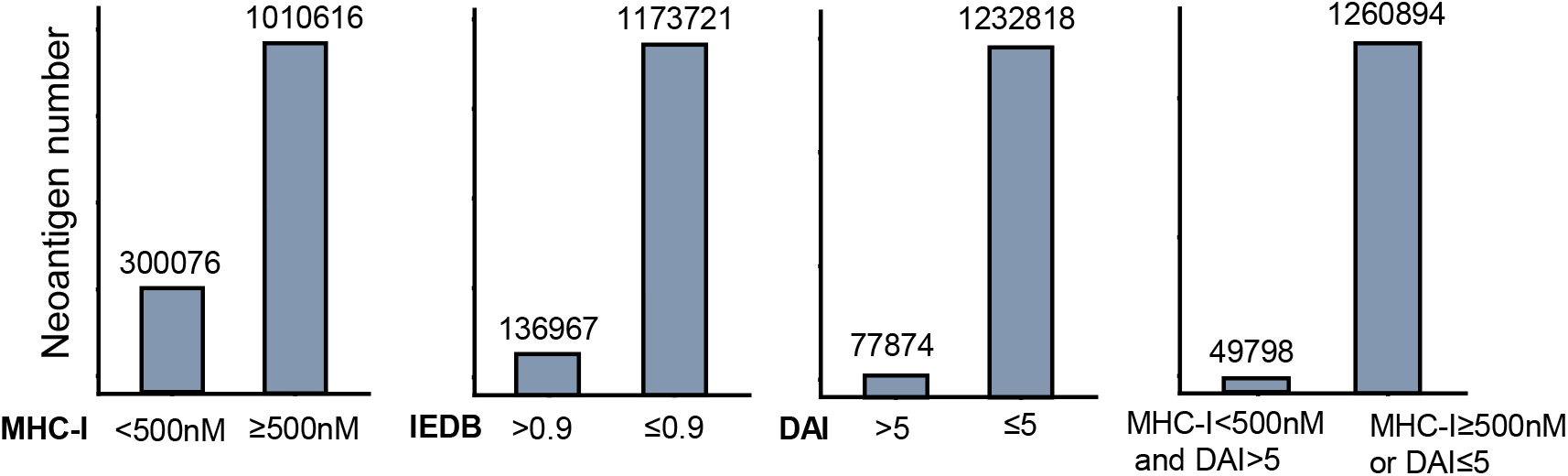
Distribution analysis of neoantigens with immunogenicity or without immunogenicity defined by four different methods.

**Extended Data Fig. 2.**
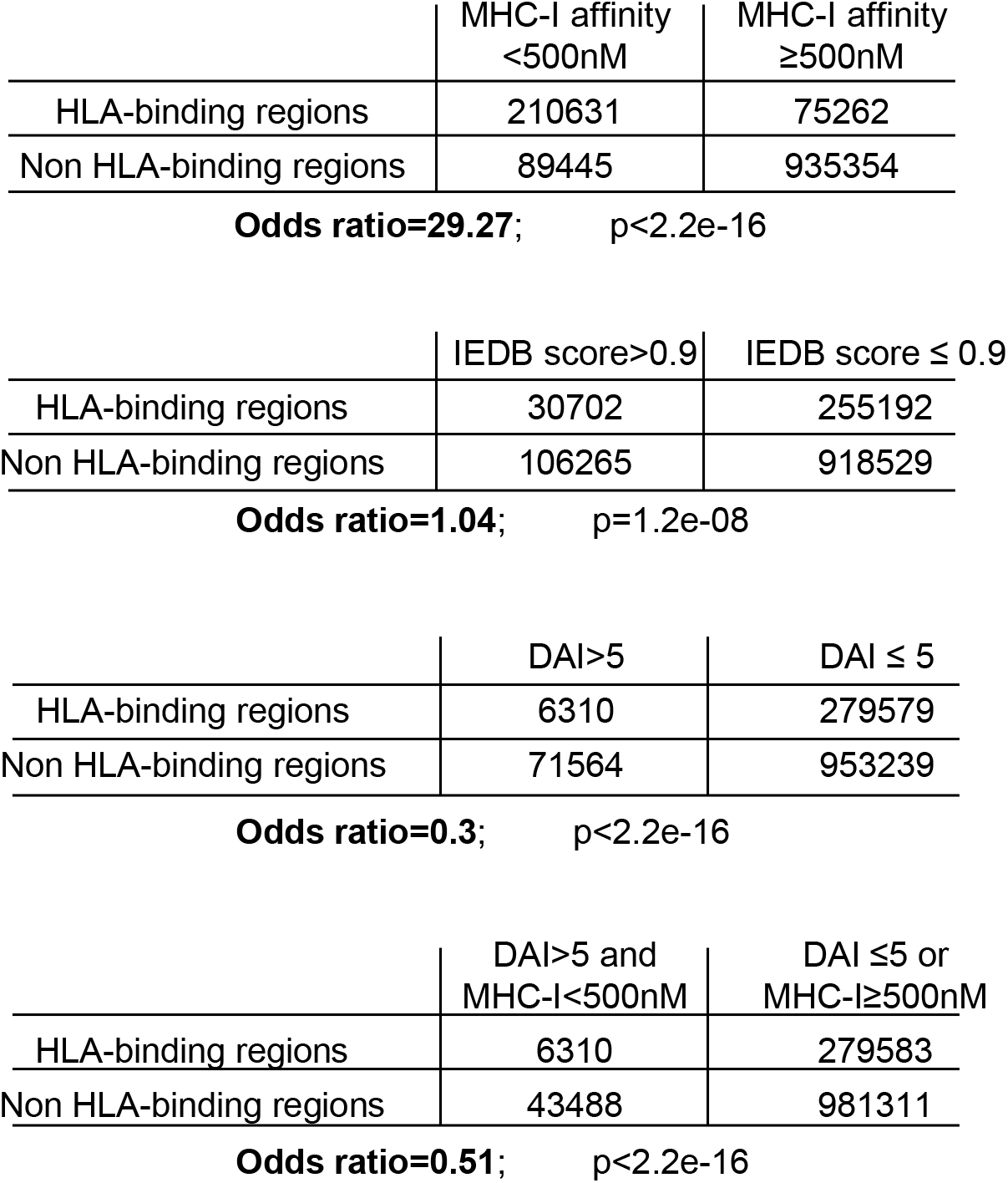
Statistic analysis for the enrichment of neoantigens with immunogenicity defined by four different methods in “HLA-binding regions”. Fisher extract test was performed and p values are reported.

## References

1. Schreiber, R.D., Old, L.J. & Smyth, M.J. Cancer immunoediting: integrating immunity’s roles in cancer suppression and promotion. Science 331, 1565–70 (2011).

2. Van den Eynden, J., Jimenez-Sanchez, A., Miller, M.L. & Larsson, E. Lack of detectable neoantigen depletion signals in the untreated cancer genome. Nat Genet 51, 1741–1748 (2019).

3. Lawrence, M.S. et al. Mutational heterogeneity in cancer and the search for new cancer-associated genes. Nature 499, 214–218 (2013).

4. Richters, M.M. et al. Best practices for bioinformatic characterization of neoantigens for clinical utility. Genome Med 11, 56 (2019).

5. Vita, R. et al. The immune epitope database (IEDB) 3.0. Nucleic Acids Res 43, D405–12 (2015).

6. Luksza, M. et al. A neoantigen fitness model predicts tumour response to checkpoint blockade immunotherapy. Nature 551, 517–520 (2017).

7. Duan, F. et al. Genomic and bioinformatic profiling of mutational neoepitopes reveals new rules to predict anticancer immunogenicity. J Exp Med 211, 2231–48 (2014).

8. Bjerregaard, A.M. et al. An Analysis of Natural T Cell Responses to Predicted Tumor Neoepitopes. Front Immunol 8, 1566 (2017).

9. Rech, A.J. et al. Tumor Immunity and Survival as a Function of Alternative Neopeptides in Human Cancer. Cancer Immunol Res 6, 276–287 (2018).

10. Richman, L.P., Vonderheide, R.H. & Rech, A.J. Neoantigen Dissimilarity to the Self-Proteome Predicts Immunogenicity and Response to Immune Checkpoint Blockade. Cell Syst 9, 375–382 e4 (2019).

11. Supek, F., Minana, B., Valcarcel, J., Gabaldon, T. & Lehner, B. Synonymous mutations frequently act as driver mutations in human cancers. Cell 156, 1324–1335 (2014).

12. Quinlan, A.R. & Hall, I.M. BEDTools: a flexible suite of utilities for comparing genomic features. Bioinformatics 26, 841–2 (2010).

